# Microinjection to deliver protein, mRNA, and DNA into zygotes of the cnidarian endosymbiosis model *Aiptasia* sp.

**DOI:** 10.1101/187278

**Authors:** Victor A. S. Jones, Madeline Bucher, Elizabeth A. Hambleton, Annika Guse

## Abstract

Reef-building corals depend on an intracellular symbiosis with photosynthetic dinoflagellates for their survival in nutrient-poor oceans. Symbionts are phagocytosed by coral larvae from the environment and transfer essential nutrients to their hosts. *Aiptasia*, a small tropical marine sea anemone, is emerging as a tractable model system for coral symbiosis; however, to date functional tools and genetic transformation are lacking. Here we have established an efficient workflow to collect *Aiptasia* eggs for *in vitro* fertilization and microinjection as the basis for experimental manipulations in the developing embryo and larvae. We demonstrate that protein, mRNA, and DNA can successfully be injected into live *Aiptasia* zygotes to label actin with recombinant Lifeact-eGFP protein; to label nuclei and cell membranes with NLS-eGFP and farnesylated mCherry translated from injected mRNA; and to transiently drive transgene expression from an *Aiptasia*-specific promoter, respectively, in embryos and larvae. These proof-of-concept approaches pave the way for future functional studies of development and symbiosis establishment in *Aiptasia*, a powerful model to unravel the molecular mechanisms underlying intracellular coral-algal symbiosis.

**Summary Statement:** Toolkit extension: development of microinjection for cellular labelling, expression of exogenous genes and live imaging in *Aiptasia*, an emerging model for intracellular coral-algal symbiosis.

## Introduction

A complex yet fundamental puzzle in cell and developmental biology is how cells from different phyla coexist and co-function in symbiosis – how do very different cells encounter each other, what molecular conversations occur to promote symbiosis establishment, and how is such a complex partnership maintained in the steady state? An ecologically crucial symbiosis is that between reef-building corals and their single-celled dinoflagellate algae symbionts, which provide photosynthetically derived nutrients underlying the productivity and health of coral reef ecosystems (Muscatine, 1990; Yellowlees, 2011). Strikingly, most corals must re-establish this vital symbiosis anew every generation in the larval or juvenile stage by taking up algal cells from the environment into the gastric cavity, after which they are phagocytosed into endodermal cells and reside as endosymbionts inside a specialized organelle, the symbiosome (van Oppen et al., 2001; Wakefield and Kempf, 2001; Rodriguez-Lanetty et al., 2009; Harii et al., 2009). Despite its importance, we know little regarding the molecular basis of the establishment of coral-algal symbiosis during development.

We are only slowly making progress towards a better understanding of symbiosis establishment, primarily due to the historical lack of tools and workable laboratory systems: reef-building corals grow slowly, are sensitive to environmental conditions, and typically sexually reproduce to spawn larvae only once annually (Babcock, et al., 1986; Technau and Steele, 2011; Harrison, 2011). Likewise, most other current cnidarian laboratory models such as *Hydra*, *Nematostella*, *Clytia*, and *Hydractinia* are not symbiotic (the exception, *Hydra viridissima*, hosts symbionts unrelated to those in corals and lacks a free-swimming larval stage). The advent of modern molecular tools has made establishing new model systems far more feasible, and we and our colleagues have developed the small marine anemone *Aiptasia* into a powerful model for molecular studies of coral-algal symbiosis (Weis et al., 2008; Goldstein and King, 2016). Housing the same symbionts as corals yet tractable in the laboratory, *Aiptasia* now has a range of key resources including a sequenced genome, transcriptomes (e.g. symbiotic *vs.* non-symbiotic), advanced microscopy, phenotypic assays, and controlled sexual reproduction in the laboratory (Sunagawa et al., 2009; Lehnert et al., 2014; Xiang et al., 2014; Baumgarten et al., 2015; Grawunder et al., 2015; Bucher et al., 2016). Just as in corals, *Aiptasia* larvae phagocytose symbionts from the environment (Hambleton et al., 2014; Wolfowicz et al., 2016).

Despite its success, *Aiptasia* has so far lacked certain tools important for a cell and developmental model system: namely, the introduction of exogenous material or the perturbation of endogenous processes. Such ability would be especially useful to study how *Aiptasia*, and by extension reef-building corals, establish symbiosis anew in the developing larval stage. To this end, microinjection of material into embryos has proven an efficient method of genetic engineering in many models, while also making possible the direct production of F0 manipulated larvae and juveniles for immediate phenotypic analysis.

There are currently no published reports of microinjection in *Aiptasia* nor any symbiotic cnidarian, with one exception of an elegant study involving the injection of morpholinos into recently spawned coral embryos (Yasuoka et al., 2016). While an important step forward, the aforementioned difficulties of efficient laboratory work in corals means that a more fruitful long-term approach would be developing these techniques in the *Aiptasia* model. We have excellent guides not only from that study but also from work in the cnidarian models *Hydra*, *Nematostella*, *Clytia*, and *Hydractinia*, in all of which successful microinjection and genetic engineering protocols are employed (Wittlieb et al., 2006; Momose et al., 2007; Renfer et al, 2009; Kunzel et al., 2010; Marlow et al., 2012; Layden et al., 2013, Ikmi et al., 2015; Artigas et al., 2017).

The development of microinjection and the subsequent introduction of foreign material would be ground-breaking for the *Aiptasia* and coral-algal symbiosis fields. It would open the door to myriad observational and functional studies, propelling the symbiosis field forward and allowing both broad approaches as well as specific hypothesis testing based on candidate genes. To this end, here we show the establishment of microinjection in the *Aiptasia* model system in a simple and robust workflow to introduce exogenous material into embryos for subsequent analysis. We describe conditions for regular gamete production as a prerequisite to efficient microinjection. We then show successful introduction of three key materials: fluorescent protein, mRNA, and DNA plasmids, with visualization of the fluorescent products in both fixed and live samples. Importantly, introduction of such exogenous material appears to have no significant effects on either development or symbiosis establishment, demonstrating the utility of these tools to study fundamental questions of development and symbiosis establishment in the *Aiptasia* system.

## Results

### Optimized spawning system and *in vitro* fertilization for zygote production

An important prerequisite for zygote microinjection is control over fertilization, which allows regular access to developmentally synchronized stages. We previously established a robust, consistent protocol for laboratory induction of *Aiptasia* spawning based on a blue light cue (simulated full moon) (Grawunder et al., 2015). Building on this, here we optimized a controlled anemone cultivation system to induce spawning in female and male anemones separately for gamete collection and *in vitro* fertilization. Staged sets of sex-segregated mature adults were induced to spawn for two consecutive months (two lunar simulations), with spawning typically three to four weeks after the start of each cue (Fig. S1A). Gametes were produced on average 2.7 times per week (n=24 weeks) (Fig. 1A). After spawning, eggs were often in a discrete patch near the female (Fig. 1B). Sperm was sometimes seen as an obvious expelled cloud or as milky water, although often it was too dilute to directly observe (data not shown).

**Figure 1:**
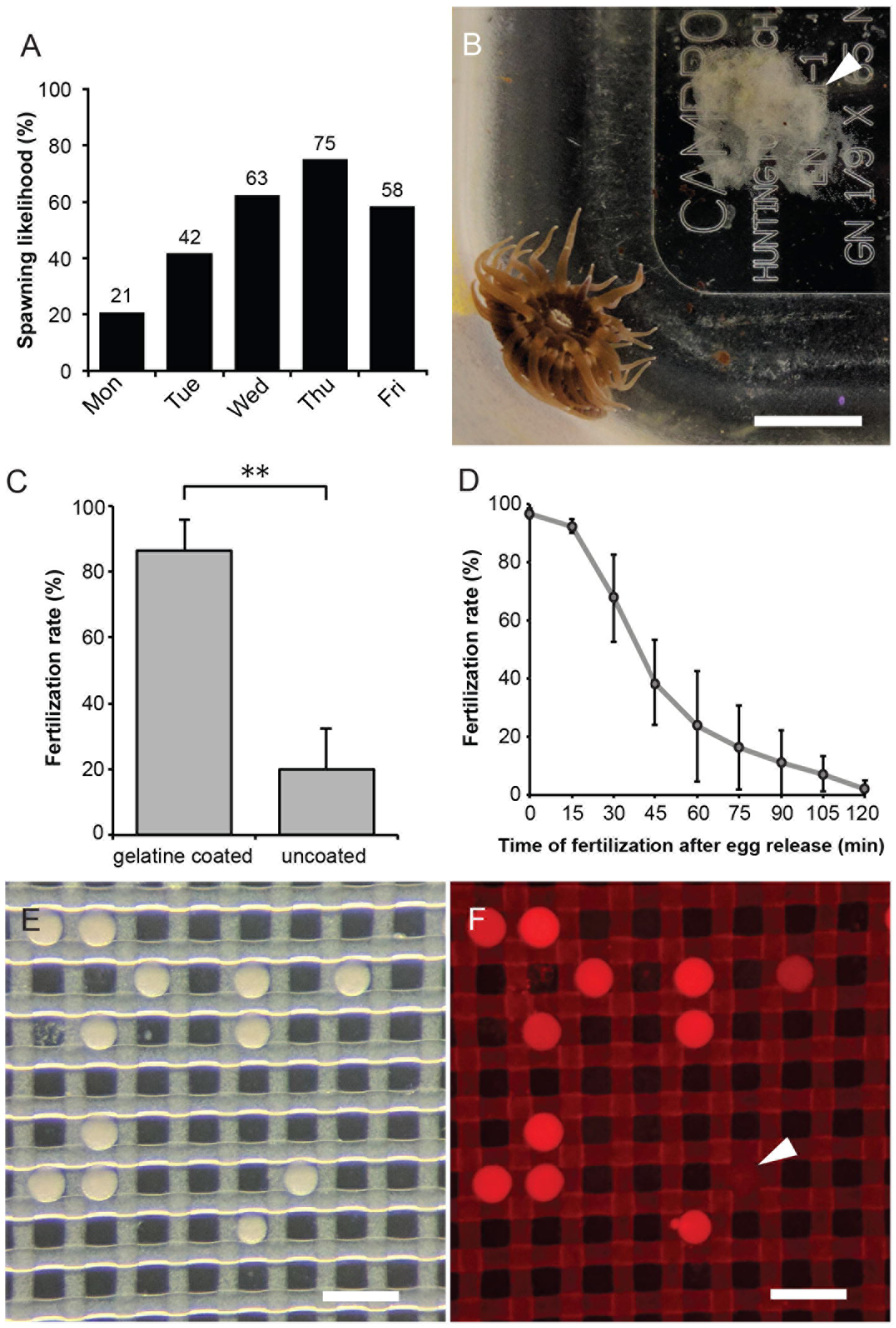
Spawning induction and *in vitro* fertilization in *Aiptasia*. (**A**) Culture and induction regime provides gametes throughout the week. Percentage of spawning events on each weekday (total from 24 weeks). (**B**) Female animal with egg patch (arrowhead). Scale = 5 mm. (**C**) Gelatin coating increases fertilization success. n=3; *p*<0.01; error bars, s.d. (**D**) Fertilization success rapidly decreases over time after spawning (n=5). Error bars, s.d. (**E**) Zygotes are transferred into a microinjection dish filled with FASW, where they sink into holes in the mesh. Scale = 200 μm. (**D**) Co-injection of dextran-AlexaFluor 594 allows identification of successfully injected zygotes. Injected zygotes are brightly fluorescent whereas uninjected zygotes appear dark (arrowhead). Scale = 200 μm.

We next assessed the efficiency of *in vitro* fertilization (IVF) of mixed spawned gametes. Several hundred eggs were gently transferred into a small plastic petri dish followed by addition of sperm-containing water. Fertilization efficiency was quantified after approximately 4-5 h, when developing zygotes could be clearly distinguished from unfertilized eggs. We found that on average, only 20% of eggs were fertilized in uncoated petri dishes; however, using dishes that were pre-coated with 0.1% gelatin in distilled water yielded mean fertilization rates of 87% (Fig. 1C). Coating dishes with 1% BSA in distilled water or agarose in filtered artificial seawater (FASW) was equally effective (data not shown). We then quantified IVF efficiency over a timecourse after spawning. On average, while more than 90% of eggs were fertilized when sperm was added within 15 min after egg release, fertilization rates fell rapidly over time, reaching 20% when sperm was added to eggs 60 min post-spawning and nearly 0% after 120 min (Fig. 1D). Thus, time is of the essence to ensure high fertilization rates.

### Establishment of an efficient *Aiptasia* zygote microinjection procedure

After undergoing IVF in coated dishes, zygotes were gently transferred to a dish with a strip of nylon mesh (80 × 80 μm) as a substrate to immobilize them for injection (Fig. S1B). With a mean diameter of 86 μm (Bucher et al., 2016), *Aiptasia* zygotes settle well into the mesh (Fig. 1E). Using a stereomicroscope set-up (Fig. S1C), material was injected to approx. a third to half of the cell diameter, corresponding to approximately 10% of the egg volume as assessed visually by the tracer dye or fluorescent protein. This fluorescence was then used to distinguish injected from non-injected zygotes (Fig. 1F). In *Aiptasia* zygotes, the first clear indication of development is the appearance of 4-cell stages approx. 90 min post-fertilization. Shortly beforehand, zygotes appear box-shaped (Fig. S1D) and we consequently stop injection; this ensures that material is delivered to cells prior to the first cleavage. Occassionally, we observed two discrete nuclei before the first obvious cleavage, likely indicating that the first nuclear division happens without cytokinesis. The next cleavage occurs approx. 20 min after the first, and embryonic development continues until the blastula stage approx. 5 h post-fertilization (hpf), at which point successfully developing embryos are easily distinguished from unfertilized eggs (arrowhead, Fig. S1E). With practice, one can inject more than 100 zygotes in the roughly 1 h window before cleavage begins.

We quantified the survival rates after 24 h of injected embryos versus those handled identically but not injected. Embryos injected with any material (i.e. protein and mRNA; see below) had a survival rate of approx. 49% (332 of 677 injected embryos). In comparision, the survival rate of control embryos was approx. 61% (306 of 501 control embryos), lower but not significantly different from the injected set (Student’s 2-tailed unpaired *t*-test, *p*-value =0.24).

### Microinjection of fluorescent protein into *Aiptasia* larvae

To visualize cell outlines in the developing larvae, we recombinantly expressed and purified Lifeact-eGFP protein (Fig. S2), which in other systems labels the actin cytoskeleton and has little to no effect on live actin cellular dynamics (Riedl et al., 2008; Sliogeryte et al., 2016). Microinjected protein instantly and ubiquitously labeled cell outlines in *Aiptasia* zygotes from the first cell divisions to 24 hours post-fertilization (hpf), especially in the ectoderm (Fig. 2A). Prominent but weaker staining was visible 48 hpf, but staining intensity decreased with larval age and was nearly undetectable 4 days post-fertilization (dpf). Consistent with the survival data above, we did not observe any appreciable defects or delays in development in injected embryos relative to uninjected controls (data not shown).

**Figure 2:**
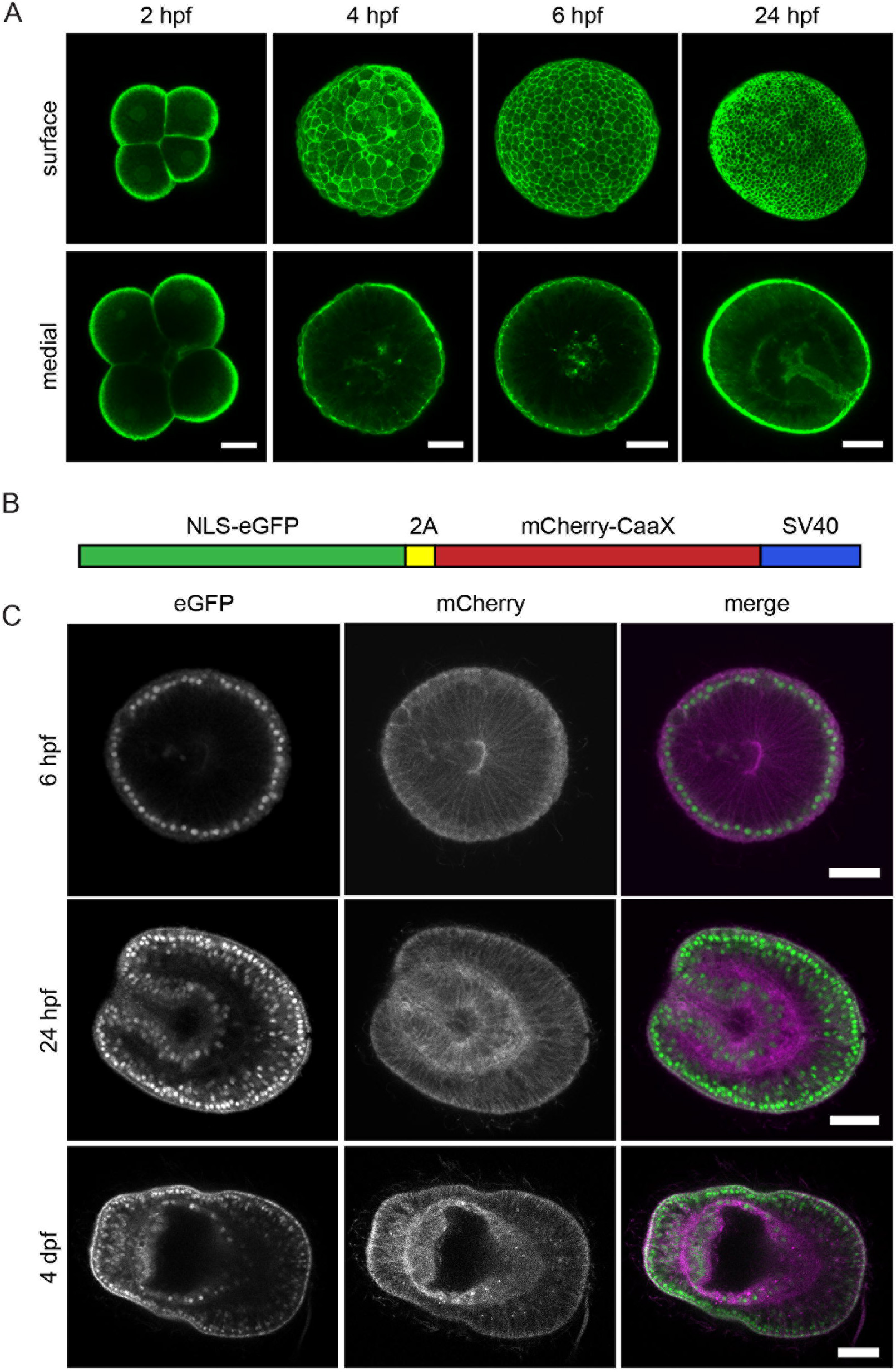
Microinjection of recombinant Lifeact-eGFP protein and mRNA to fluorescently label nuclei and cell outlines. (**A**) Zygotes injected with ~3.4 mg/ml Lifeact-eGFP protein were fixed 2, 4, 6 and 24 hpf. Maximum projections of 5 z-planes near the surface or centre of the embryo are shown. Scale = 25 μm. (**B**) Schematic representation of bicistronic *in vitro* transcribed *NLS-eGFP-V2A-mCherry-CaaX* mRNA: eGFP with a nuclear localization signal (NLS) is coupled to mCherry with a C-terminal CaaX box for farnesylation and insertion into the membrane. The fluorophores are separated by the self-cleaving V2A peptide. (**C**) mRNA-injected embryos were fixed 6 or 24 hpf or 4 days post-fertilization (dpf). Left panels show eGFP-labeled nuclei, middle panels show mCherry-labeled membranes, and the right panel shows the merged images (eGFP green; mCherry magenta). Scale = 25 μm.

### Microinjection and *in vivo* translation of exogenous mRNA in *Aiptasia* larvae

To visualize cell outlines on a longer timescale, as well as to test whether exogenous mRNA is efficiently translated in *Aiptasia* zygotes, we next injected *in vitro* transcribed bicistronic mRNA from a plasmid encoding eGFP with a nuclear localization signal (NLS-eGFP) and mCherry with a farnesylation signal (mCherry-CaaX) separated by the self-cleaving V2A peptide (Fig. 2B). This was previously shown to simultaneously label cell membranes and nuclei in developing embryos of the anemone *Nematostella* (Ikmi et al., 2014). Injected mRNA was translated robustly as indicated by strong, homogeneous, and long-lasting fluorescent labeling of the cellular structures (Fig. 2C). Labeled membranes and nuclei were detected as early as 4 hpf. At 6 hpf (blastula stage), signal intensity had further increased and remained strong for 1-2 dpf, with signal still visible at 4 dpf (Fig. 2C).

### Symbiosis establishment and live imaging of mRNA-injected larvae

Importantly for the applicability of this technique to the study of symbiosis, mRNA injection did not substantially affect symbiont uptake by *Aiptasia* larvae. Using our standard assay (Bucher et al., 2016), we exposed injected and control larvae to a compatible symbiont strain and analyzed symbiosis establishment. Injected larvae appeared developmentally normal and took up symbionts from the environment; imaging by confocal microscopy showed phagocytosed algae within the host endoderm that could be clearly distinguished from those in the gastric cavity (Fig. 3A). In four experiments, each with matched injected and control larvae, infection efficiencies did not differ significantly between mRNA-injected larvae (47%) and non-injected controls (66%) (Student’s 2-tailed unpaired *t*-test, *p*-value = 0.14) (Fig. 3B). Likewise, the average number of algal cells that each infected larva contained was similar (3.8 per injected larva vs. 3.7 per control larva; Student’s 2-tailed unpaired *t*-test, *p*-value = 0.37) (Fig. 3C). Excitingly, we observed mCherry-labeling of symbiosome membranes surrounding internalized algae; this was apparent even in cases where the larger host cell membrane labeling was difficult to distinguish (Fig. 3D). The intensity of symbiosome labeling and algal red autofluorescence varied between symbionts (Fig. S3A).

**Figure 3:**
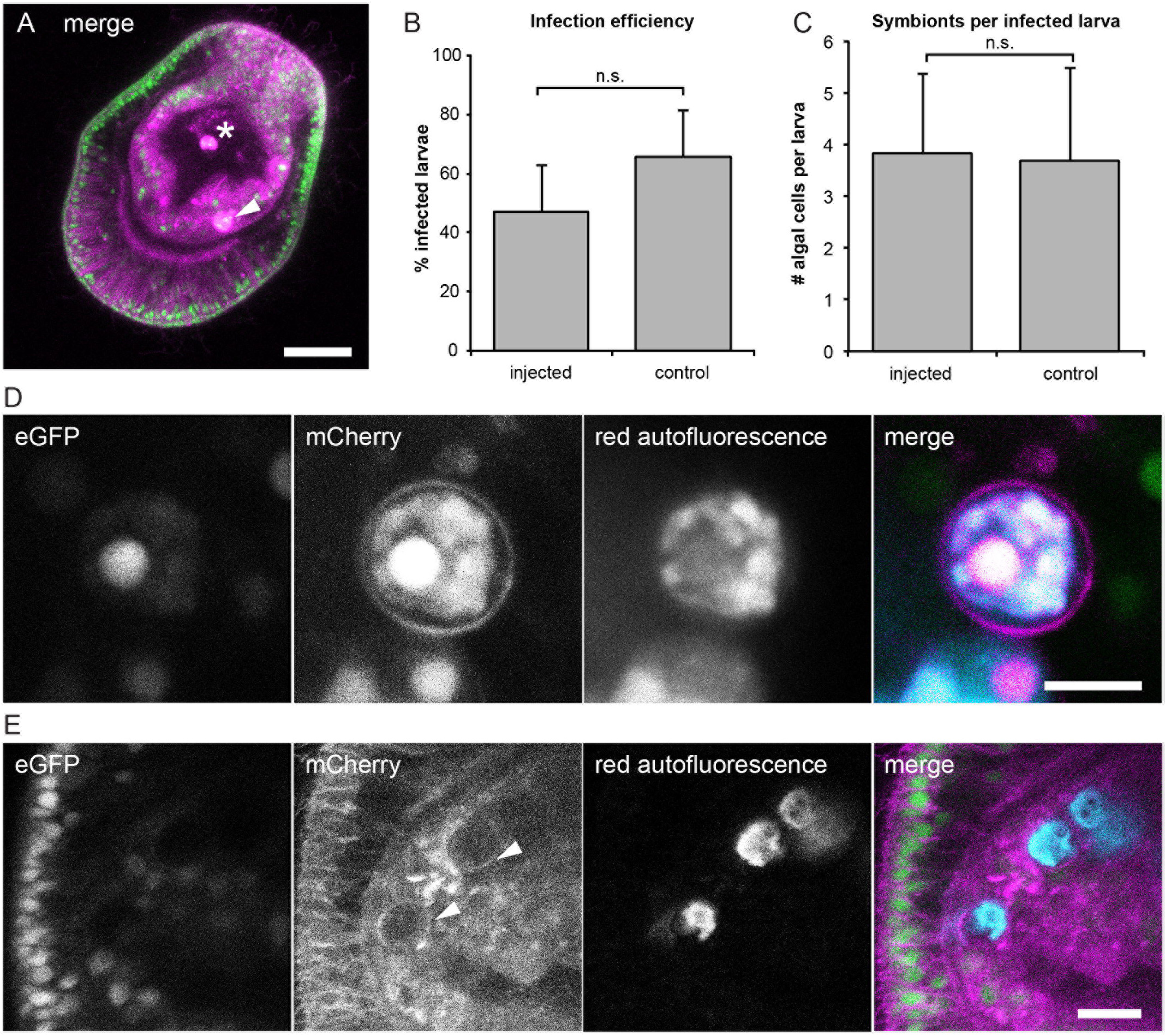
Symbiosis establishment and live imaging in microinjected larvae. (A) Larvae expressing the injected *NLS-eGFP-V2A-mCherry-CaaX* mRNA and containing acquired symbiont cells (brightly autofluorescent in magenta mCherry channel). Symbionts within endodermal tissue (arrowhead) can be distinguished from those in the gastric cavity (asterisk). Larvae were incubated with *Symbiodinium* strain SSB01 at a concentration of 100,000 cells per ml for 3 days. Scale = 25 μm. (**B**) Infection efficiencies and (**C**) average number of algae per larva do not significantly differ in mRNA injected and control larvae. (**D**) Symbionts autofluoresce in the mCherry channel (magenta) and the red channel (cyan). The mCherry-labeled symbiosome membrane can be clearly seen surrounding the internalized symbiont. Larva were injected with *NLS-eGFP-2A-mCherry-CaaX* mRNA and incubated for 1 day with *Symbiodinium* strain SSB01, before fixation at 3 dpf. Scale = 5 μm. (**E**) Symbionts and symbiosome membranes (arrowheads) can be imaged *in vivo.* Larva injected with *NLS-eGFP-2A-mCherry-CaaX* mRNA and incubated for 1 day with *Symbiodinium* strain SSB01, before embedding in agarose and imaging at 3 dpf. Scale = 10 μm.

In order to observe symbionts in living hosts, we immobilized larvae injected with *NLS-eGFP-V2A-mCherry-CaaX* by embedding them in agarose; this prevented larvae from swimming, but their rotation caused by ciliary beating continued (Movie S1). Using this technique, nuclei, cell membranes, and the symbiosome could be imaged live (Fig. 3E), at a rate of 1 frame per 2.5 sec (Movie S2), which would allow cellular events to be followed in real time. Embedded larvae survived and could be imaged for several hours (Fig. S3B).

### Microinjection of DNA in *Aiptasia* larvae

With the goal of achieving gene expression from exogenous DNA and ultimately establishing stable transgenesis, we generated plasmids in which the *NLS-eGFP-2A-mCherry-CaaX* reporter was driven by the promoter of different actin genes cloned from *Aiptasia* (Fig. 4A,B). In each case, the cassette was flanked by recognition sites for the meganuclease I-SceI (Fig. 4A); co-injection of I-SceI with plasmids containing these sites strongly increased genomic integration in other species, e.g. medaka and *Nematostella* (Grabher and Wittbrodt 2007; Renfer and Technau 2017). We identified six actin genes from the *Aiptasia* genome (Baumgarten et al., 2015), and used larvae transcriptome data (Wolfowicz et al., 2016) to select the four highest expressed to clone and use for injection (Fig. 4B). The percentage of larvae displaying reporter expression at 10 hpf was determined, and was clearly the highest for one promoter (Accession # XM_021049442.1, 63%, n=97 larvae, Fig. 4B). Reporter expression was mosaic in these larvae, with cell patches of varying sizes displaying GFP-labeled nuclei and mCherry-labeled membranes (Fig. 4C). The three other *Aiptasia* promoters tested (Fig. 4B) yielded low proportions of larvae with individual fluorescent cells rather than whole fluorescent patches. For larvae expressing reporters driven by the XM_021049442.1 promoter, we did not determine whether the transgene was stably integrated into the genome, but the successful *in vivo* expression allows future work to establish *Aiptasia* transgenesis.

**Figure 4:**
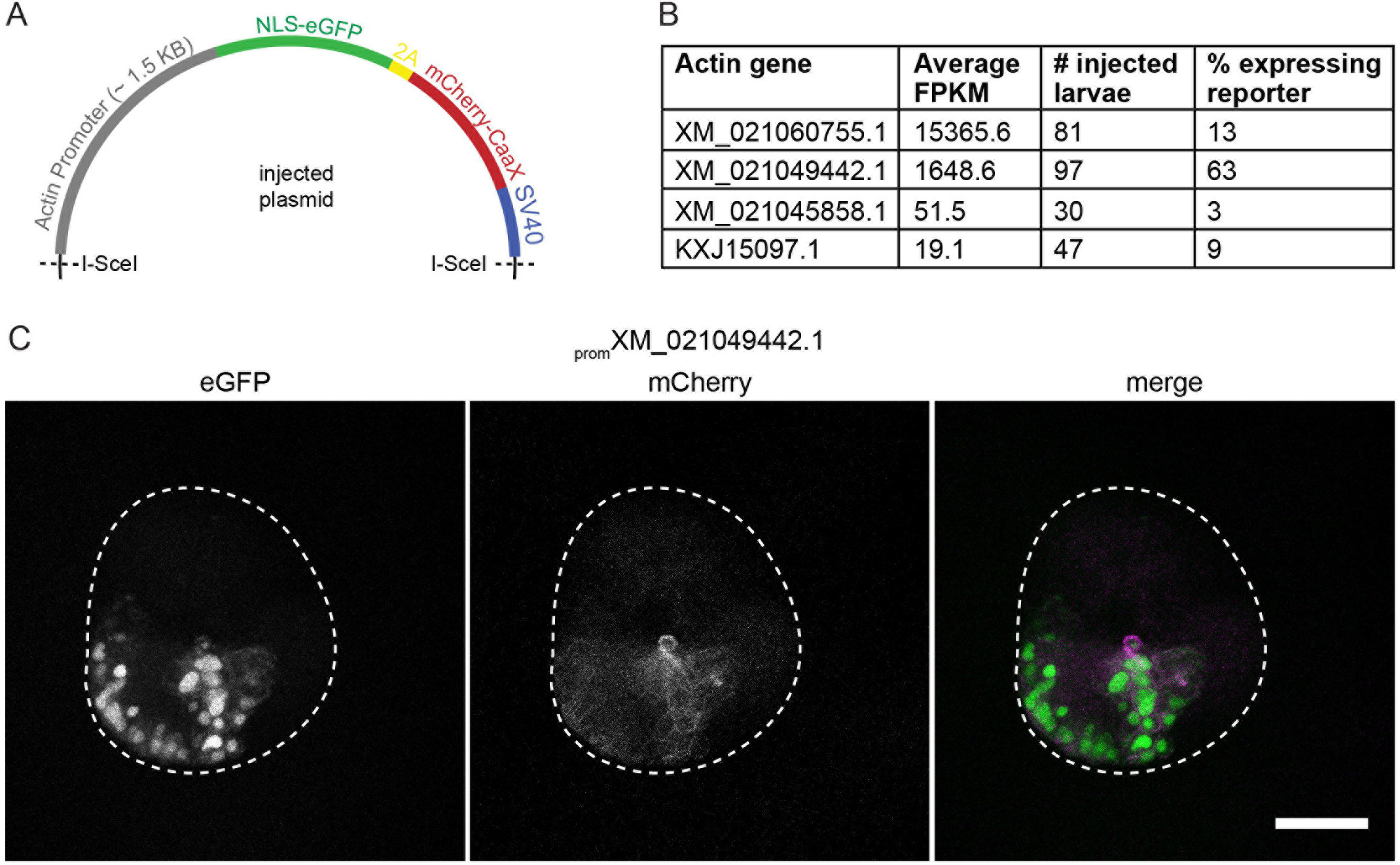
Microinjection of DNA into *Aiptasia* zygotes. **(A)** Schematic of injected plasmids: the promoter of an *Aiptasia* actin gene drives expression of the *NLS-eGFP-V2A-mCherry-CaaX* reporter with a SV40 termination sequence. Meganuclease I-SceI recognition sites are indicated. (**B**) The promoter of actin gene XM_021049442.1 drives reporter expression in the majority of larvae coinjected with plasmid and the meganuclease I-SceI. For each promoter tested, the expression level (FPKM) (larvae transcriptomes, Wolfowicz et al., 2016), the number of larvae injected and analyzed, and the percentage of larvae in which GFP signal could be detected are given. (**C**) Coinjection of I-SceI and the _*prom*_ XM_021049442.1: *NLS-eGFP-V2A-mCherry-CAAX:SV40* plasmid causes strong, mosaic expression of the transgene, labeling nuclei (green) and membranes (magenta). 10 hpf, scale = 25 m.

## Discussion

Here we establish a workflow to successfully introduce exogenous protein, mRNA, and DNA via microinjection into *Aiptasia* zygotes and, critically, we demonstrate that such manipulation has no significant effects on development or symbiosis establishment in *Aiptasia* larvae. This progress propels the *Aiptasia* model system forward in answering fundamental questions regarding development and concurrent establishment of coral-algal symbiosis. While immediately permitting a range of observational and functional assays, it also opens the door to CRISPR-Cas9-induced gene editing and the production of stable transgenic lines. Finally, this work holds broader implications for comparative developmental biology and emerging technologies in the current bloom of new model systems across the life sciences.

This workflow immediately permits in the *Aiptasia* larval system many observational and functional assays that bring us closer to understanding larval development and symbiosis establishment. The small transparent larvae of *Aiptasia* are amenable to microscopy, which was until now used to assay fixed specimens (Hambleton et al., 2014; Bucher et al., 2016). Thus, the investigation of development and the symbiosis establishment process was necessarily limited to “snapshots”; now, live imaging permits observation of development and symbiosis dynamics in real time, with injected genetic material or even commercially available dyes to label conserved cellular structures.

To study developmental processes, larvae with labeled cell outlines (Figs 3, 4) and live imaging can be used to characterize, for example, gastrulation, tissue differentiation, and cell division and migration. Injected recombinant Lifeact-eGFP protein instantly labels all cells in developing zygotes, allowing studies of very early embryonic development. Similarly, injection of *NLS-eGFP-V2A-mCherry-CaaX* mRNA allows monitoring of cellular dynamics once translation has started. Early and homogenous expression of exogenous mRNA also allows the manipulation of developmental genes, such as those involved in patterning and axis establishment, to monitor dynamics or create overexpression phenotypes.

To study symbiosis establishment in *Aiptasia* larvae, key goals are to test candidates for their roles in symbiosis as well as to observe symbiont phagocytosis and proliferation. Importantly, we can immediately tackle the first goal by using injected mRNA. For instance, we can now express proteins fused to fluorescent tags to track their co-localization with symbionts or other proteins, or analyze the effects of the over-expression phenotypes of symbiosis-specific genes on symbiosis establishment. The constructs reported here have proven limited for observing symbiont phagocytosis and proliferation with cellular resolution; both the directly injected Lifeact-eGFP protein and exogenously expressed farnesylated mCherry lose resolution in the endoderms of older larvae (Figs 3,4). This phenomenon was also reported in *Nematostella*, where signal from some mRNA lasts for two months after injection yet others are rapidly lost (DuBuc et al., 2014). There is however overlap between signal and symbiosis, as *Aiptasia* larvae acquire symbionts at and after 2 dpf, and signal of the mRNA fades several days later. Furthermore, the unexpectedly strong labeling of symbiosome membranes by farnesylated mCherry, even in older larvae, is a great advantage to studies of intracellular symbiosis dynamics.

We have made progress towards the goal of generating stable transgenic lines by demonstrating expression of an injected DNA construct with a fluorescent reporter driven by an *Aiptasia* endogenous actin promoter. To date we have only achieved transient and mosaic expression, but this is a key first step for future transgenesis attempts. Currently, a drawback in the *Aiptasia* system is that metamorphosis and settlement of larvae into adults cannot yet be accomplished in the laboratory; multiple groups are actively working towards identifying the cue to induce closure of the life cycle. Nevertheless, the identification of a functional *Aiptasia* promoter may encourage testing alternative approaches to generating stable transgenic lines. For example, constructs could be delivered via gene bombardment (Böttger *et al,.* 2002) or electroporation (Bosch *et al.*, 2002; Watanabe *et al.*, 2014) to *Aiptasia* adults, which rapidly reproduce asexually and have a high regenerative capability. Dissection of mosaic adults and subsequent regeneration would create lines with germline transgene transmission.

The *Aiptasia* field, and by extension the coral-algal symbiosis field, acutely requires tools to translate knowledge on molecular players into a mechanistic understanding at the functional level. In addition to the gain-of-function possibilities outlined above, the ability to deliver materials by microinjection facilitates the next major leap forward for the *Aiptasia* model: gene editing by CRISPR-Cas9. Such gene editing would allow, for the first time, knock-out of candidate genes implicated in symbiosis establishment to unequivocally and functionally demonstrate their role in symbiosis. It is apparent that when developing a new model, not all techniques can be established at once. While some techniques must await, for example, the closure of the *Aiptasia* life cycle, other techniques can be employed immediately for urgent questions. Here we show that *Aiptasia* zygotes can be injected in sufficient numbers and the larvae used for symbiosis studies in the F0 generation.

Beyond the *Aiptasia* system, this work holds broader implications for comparative developmental biology and other emerging model systems. The studies of embryonic development in *Aiptasia* discussed above would complement those in other systems to dissect the evolution of fundamental developmental processes, such as gastrulation. As the sister group to bilaterians, cnidarians are important “evo-devo” models to infer evolutionary conservation and divergence of development (reviewed in Layden et al., 2016). The advent of modern research tools has led to rapid advances in emerging models, as new avenues are opened to study previously intractable cell biological questions (Cook et al., 2015; Goldstein and King, 2016). The *Aiptasia* system is currently undergoing this transition: a wealth of resources has been recently and rapidly built, but missing was the transformative power of manipulation via introduction of exogenous material or targeted functional analysis. Our progress on this front was inspired by successful techniques in other model systems, and we hope this work in turn provides helpful “lessons learned” for other emerging models. With the rapid development of myriad new model systems, we are in the midst of an exciting time for major discoveries in underexplored cell biological phenomena.

## Material and Methods

### Anemone cultivation for spawning induction

Individuals to be used for gamete production were produced asexually by pedal laceration from adult animals in master stock tanks of either male CC7 (Sunagawa et al., 2009) or female F003 (Grawunder et al., 2015) clonal lines. To raise them to sexual maturity, 12-14 medium-sized animals of each line (with an oral disc diameter of 4-5 mm for F003 and 5-6 mm for CC7) were maintained for 6-7 months as previously described (Grawunder et al., 2015), in covered, food-grade plastic tanks in a volume of approximately 1.6 l artificial sea water (ASW). Briefly, tanks were kept at 26°C with a 12L:12D photoperiod from 8 am to 8 pm to allow observation and maintenance during the daytime. Animals were fed five times a week with fresh *Artemia* nauplii. Three times per week, the surfaces of the tanks were cleaned with cotton swabs and ASW was exchanged. During this period, anemones of both lines grew to an oral disc diameter of 11-12 mm.

### Spawning induction of *Aiptasia*

To prepare sexually segregated tanks of *Aiptasia* for gamete release, 3-5 mature animals of either CC7 or F003 were transferred into 300 ml ASW in smaller tanks (#92CW, Cambro, USA) one week before spawning induction to allow acclimatization to the new tank. Tanks were fed and cleaned as described above. Tanks were then kept at 29°C in Aqualytic Incubators (Model TC 135 S, Liebherr, Germany) equipped with white LEDs (SolarStinger Sunstrip “Marine”, # 00010446, Econlux, Germany) at an intensity of 23-30 μmol m^−2^ s^−1^ on a 12L:12D photoperiod with darkness from 4 pm to 4 am, thereby adjusting the animals’ diurnal rhythms to allow gamete collection during working hours. To induce gamete release following a simulated full moon cue, animals were exposed to blue light LEDs (SolarStinger Sunstrip “Deepblue”, # 00010447, Econlux, Germany) at 15-20 μmol m^−2^ s^−1^ for the entire dark phase on days 1 to 5 of a 28-day cycle (Grawunder et al., 2015). Each set of tanks was kept for two simulated lunar cycles, and sets were staggered such that the second month of one tank set overlapped with the first month of the next tank set. As a result, there was always one tank set in its first month and one in its second. Tanks were examined for the presence of eggs or sperm using a Leica S8APO stereoscope Monday through Friday from days 13 to 24 of each cycle. Spawning was checked from 9am - 10am, approx. 5-6 h after the onset of darkness.

### *In vitro* fertilization (IVF) of gametes

Eggs from F003 (female-only) tanks were transferred gently with a plastic transfer pipette into a small plastic petri dish (60 mm × 15 mm) in a volume of approx. 5-10 ml. On occasion, spawned eggs float instead of forming discrete patches; we had previously tried to concentrate these using small 40 μm filters, but the handling substantially reduced the number of normally developing embryos. We therefore used only spawning events that resulted in egg patches for microinjection (Fig. 1A). To fertilize the eggs, approx. 3-7 ml water from several induced CC7 (male-only) tanks was added to dish to maximize the chances of sperm presence and fertilization. Sperm were sometimes observed either in an obvious expelled cloud or as milky tank water, yet even when too dilute to be detected via stereoscope, they were nevertheless often present as seen in the generally high fertilization rates (also confirmed with DIC microscopy and/or Hoechst nuclear staining [data not shown]).

Noticing low fertilization rates in preliminary experiments, we compared IVF efficiency of gametes mixed in uncoated petri dishes to those in dishes pre-coated with gelatin. To coat, a 0.1% solution of gelatin (# G1393, Sigma-Aldrich, Germany) in distilled water was poured into the dishes, incubated at room temperature (RT) for 5 min, removed, and then the dishes were air-dried at RT overnight and stored at RT until further use. Eggs were added to the dishes as described above, and sperm-containing water was then added at the indicated time-points in Fig. 1D. Based on the outcome of these comparisons (Figs 1C,D), eggs for microinjection were afterwards always prepared in a 0.1% gelatin-coated dish and mixed with water from CC7-containing tanks, as described above, as soon as possible after gamete release. The eggs and sperm were incubated together at RT for approx. 10 min to allow fertilization to occur while preparing the equipment and solutions for microinjection.

### Recombinant expression of Lifeact-eGFP protein in bacteria

The Lifeact-eGFP sequence with a C-terminal 6His-tag was PCR amplified from the plasmid described by Riedl et al. (2008) and subsequently cloned into the pET21a vector using classical restriction-based cloning methods. For expression in *E.coli*, cells of strain BL21 were transformed with the plasmid via chemical transformation. A single colony of transformants was picked and grown in 1 l of LB medium to an OD_600_ of 0.6, at which time protein expression was induced by adding IPTG to a final concentration of 1 mM. Bacterial cells were recovered by centrifugation at 2,000 *xg* for 20 min, washed with 50 ml PBS, and re-centrifuged exactly as above. The pellet was then frozen in liquid nitrogen, stored at −80°C, and quick-thawed at 37°C the following day. 40 ml lysis buffer (50 mM NaPO_4_ pH 8.0, 0.5 M NaCl, 0.5% glycerol, 0.5% Tween-20, 10 mM imidazole, 1 mg/ml lysozyme) was added to each pellet and mixed to start cell lysis. After stirring for 15 min at 4°C, the lysate was sonicated on ice with several pulses (Output Control 1.2, Duty Cycle 40%, 15-30 s total) until no longer viscous. To clarify the lysate, it was spun at 37,000 *xg* for 1 h at 4°C. The supernatant was then mixed with 2 ml pre-washed Ni-NTA agarose beads (# 31105, Cube Biotech, Germany) and protein allowed to bind to the beads by rotating 2 h at 4°C. Beads with bound protein were washed three times with 20 ml wash buffer 1 (50 mM NaPO_4_ pH 8.0, 250 mM NaCl, 0.05% Tween-20, 20 mM imidazole), once with 20 ml wash buffer 2 (50 mM NaPO_4_ pH 6.0, 250 mM NaCl, 0.05% Tween-20, 20 mM imidazole) and twice 20 ml modified wash buffer 1 (50 mM NaPO_4_ pH 8.0, 250 mM NaCl, 20 mM imidazole). Between each wash, beads were pelleted by centrifuging for 3 min at 2,800 *xg* at 4°C. Beads were carefully transferred into a poly-prep chromatography column (# 7311550, BioRad Laboratories) and the protein was eluted with sequential applications of 500 μl elution buffer (50 mM NaPO_4_ pH 8.0, 150 mM NaCl, 250 mM imidazole, 5% glycerol). Collected protein fractions were analyzed using SDS-PAGE. Fractions with the highest protein content were pooled and dialyzed against 1x PBS. Protein concentration was determined via Bradford assay and absorbance measurements at A280 using a Nanodrop 1000 (Thermo Scientific). Protein aliquots were flash-frozen in liquid nitrogen and stored at −80°C until further use.

### Synthesis of mRNA for microinjection

The *NLS-eGFP-V2A-mCherry-CaaX* construct was transcribed from the pSYC-97 vector, a kind gift from Aissam Ikmi (Kim et al., 2011; Ikmi et al., 2014). mRNA was synthesized using the mMESSAGE mMachine SP6 kit (# AM1340, Thermo Fisher Scientific) and purified with the RNeasy Mini Kit (# 74104, Qiagen), according to the instructions in both kits. The quality and concentration of the mRNA was assessed on a 1% agarose gel and with a Nanodrop 1000 (Thermo Fisher Scientific). The mRNA was then diluted to 600 ng/μl with RNase-free water and single-use aliquots of 2 μl each were stored at −80°C until use.

### Cloning of DNA plasmid for microinjection

The plasmids used in this study were derived from that reported by Renfer et al. (2010). The *mCherry* CDS in the original plasmid was replaced through classical restriction-enzyme cloning with the *NLS-EGFP-V2A-mCherry-CAAX-SV40* reporter cassette amplified from the pSYC-97 vector (Kim et al., 2011). Four actin genes were selected from the six *Aiptasia* genomic actin loci using normalized expression levels (Fragments Per Kilobase of transcript per Million mapped reads; FPKM) from previously published transcriptomes (Wolfowicz et al., 2016); FPKM values of each gene model were averaged across four assemblies (two aposymbiotic and two symbiotic replicates) to generate a mean FPKM reflecting global expression in larvae. The *MyHC1* promoter was replaced through classical restriction-enzyme cloning with 1.5 kb upstream of the start codon of each of four *Aiptasia* actin genes (primers and related information in Supp. Table S1). *Aiptasia* genomic DNA (gDNA) was extracted from individual small symbiotic adults of clonal line CC7 using the DNeasy Blood & Tissue Kit (#69504, Qiagen) according to the manufacturer’s instructions, and PCR reactions to amplify the regions were each 50 μl containing 2U Phusion polymerase, 0.1 μM of each primer, 200 μM dNTPs, 1X Phusion HF Buffer (#B0518S, NEB), and 150 ng template gDNA. Amplification conditions were as follows: initial denaturation at 98°C for 2 min; 30 cycles of denaturation at 98°C for 15 s, annealing at 60°C for 20 s, extension at 72°C for 2 min; final extension at 70°C for 10 min.

### Solutions of protein, mRNA, and DNA for microinjection

For Lifeact-eGFP protein, an aliquot was quick-thawed and injected at a concentration of ~3.4 mg/ml, using the green fluorescence of the protein itself as an injection tracer. For mRNA, an aliquot of 600 ng/μl was thawed and mixed 1:1 with 0.5% phenol red in RNAse-free water as an injection tracer, for a final concentration of 300 ng/μl mRNA. For DNA plasmids, a 2 μl aliquot of DNA at 250 ng/μl was quick-thawed and mixed with 2.1 μl of either Phenol Red or Alexa-594 injection tracer (see below), 0.5 μl CutSmart Buffer (# B7204S, NEB), and 0.4 μl of the meganuclease I-SceI at 5 Units/μl (# R0694S, NEB), for a final concentration of 250 ng/μl DNA and 0.4 Units/μl enzyme. The mixture was incubated at 37°C for 10-20 min while the microinjection apparatus was assembled. Phenol red, a pH indicator that appears yellow when intracellular but red in seawater, provides a clear indication of when injection has been successful and, crucially, does not interfere with later fluorescence. Its disadvantage is that it quickly dissipates within zygotes, requiring transfer to a separate dish very soon after injection while the tracer can still be seen, interrupting the work. A fluorescent tracer is preferable when possible because it remains visible for longer and facilitates later selection of injected zygotes. We therefore use the fluorescent tracer 10,000 MW dextran coupled to Alexa-594 dye (# D22913, Invitrogen) at a final injection concentration of 0.2-0.5 μg/μl (Fig. 1F).

### Microinjection

Following Wessel et al., 2010, the injection dish was prepared ahead of time by affixing a strip of 80 × 80 μm nylon mesh (# SW10080.010.010, Meerwassershop (www. Meerwassershop.de), Germany) to the lower inside surface of a small petri dish lid (35 × 10 mm), using a line of silicon grease around the edges of the mesh (Fig. S1B). FASW was then added to cover the bottom entirely. Zygotes were concentrated in the center of the fertilization dish by gentle rotation, and slowly taken up with a 10 μl pipette with a plastic disposable tip. The zygotes were gently ejected into the injection dish under water, so that they fell sparsely in a stripe onto the mesh and settled into the holes. We noted that zygotes as well as developing embryos are sensitive and should be pipetted as gently as possible to avoid developmental defects.

Injection solution was loaded into a Femtotip needle (# 5242952008, Eppendorf) using Microloader tips (# 5242956003, Eppendorf) and allowed to run into the tip by gravity flow. Microinjections were performed with a manual micromanipulator (U-31CF micromanipulator [Narishige, Japan] with a standard universal capillary holder [# 920007392, Eppendorf] and Femtotip adapter [# 5176190002, Eppendorf]) attached to a FemtoJet 4i microinjector (# Num. 5252000013, Eppendorf). The angle of the needle holder to the bench-top was approximately 55°. Either the “constant flow” or the “injection” setting on the microinjector was used to deliver the solution into the cell. Solution flow and approx. injected amount (roughly 1/3 cell diameter or approx. 10% of cell volume) was visually assessed during the session and the pressure adjusted as necessary; during the session, the needle tip may need to be broken to remove blockages and then the injection pressure recalibrated. Microinjections were conducted either on a Nikon SMZ18 stereoscope or a Leica MSV269 stereoscope using a 0.5x objective or 1x objective, respectively. Injection was conducted either under white illumination to visualize zygotes and phenol red tracer, or, when applicable, indirect white illumination together with additional epifluorescent illumination and filters to visualize protein/tracer fluorescence. We injected by keeping the dish stationary and moving the needle between zygotes and along its own axis to enter each zygote (Fig. S1C).

Zygotes were injected for approx. 60 min, corresponding to until approx. 90 min after fertilization, at which time the first cleavages began. Zygotes injected using the tracer phenol red were transferred immediately after injection (before the tracer disappeared) by aspirating them individually into the well of a 6-well culture plate filled with approximately 5 ml of FASW using a P10 pipette. Zygotes injected with Lifeact-eGFP or other fluorescent tracer could be distinguished from non-injected zygotes and transferred at the end of the injection session. Eggs that were uninjected but otherwise handled identically in all steps, including addition to and removal from the injection dish, served as controls. Plates with microinjected embryos were kept in the dark at 29°C to develop.

### Assay of symbiosis establishment in larvae

Infection of injected or control larvae with *Symbiodinium* strain SSB01 (Xiang et al., 2013) was performed as described previously (Bucher et al., 2016). Briefly, larvae were kept in 5 ml of FASW in a 6-well plate, and algae were added to a final concentration of 100,000 algal cells/ml and mixed gently with a pipette. Algae and larvae were co-cultivated for 1 or 3 days, after which larvae were fixed, mounted on slides, and assessed by epifluorescence microscopy, as described below. Infection rates were determined by counting the number of larvae containing algae (percent infected) and the number of algal cells that each infected larva contained, as described previously (Bucher et al., 2016).

### Microscopy and image analysis

Transmitted light and fluorescence images of live zygotes in Fig. 1 and Supp. Fig. 1 were acquired using a Nikon SMZ18 binocular microscope. For imaging of fixed samples in Figs. 2,3,4, larvae were incubated for 15 min in 3.7% formaldehyde in FASW, washed three times in PBS containing 0.2% Triton X-100, and then mounted in 50% glycerol in PBS. For live imaging in Figs 1E, S3, and Supp. Movie 1, samples were embedded in agarose to restrict movement with the following method. Each imaging chamber was a small petri dish with a drilled hole in the bottom (diameter = 1.5 cm), which was then sealed by a coverslip glued to the exterior. Larvae were collected into a volume of 5-10 μl FASW and gently transferred to the dish bottom in the middle of the coverslip. 25 μl of liquid 1.8% low-melt agarose (Sigma-Aldrich, # A9414) in FASW at 37°C was added to the larvae on the coverslip and briefly mixed. A small round coverslip (Electron Microscopy Sciences, # 72230-01) was immediately placed on top and very slightly pressed. 5 ml of FASW was then added and the petri dish sealed to prevent evaporation. Images in Figure 2 were acquired using a Nikon A1 confocal microscope with a Nikon Plan Fluor 60x water immersion objective and Nikon Elements software. Images in Fig. 3 and 4 were acquired with a Leica SP8 confocal laser scanning microscope and a 63x/1.30 glycerol immersion lens using Leica LAS X software. Fluorophores were excited with 488 nm and 594 nm laser lines. GFP was detected at 493-571 nm, mCherry at 599-650 nm, and algal red autofluorescence at 656-692 nm. All image processing was performed using Fiji (Schindelin et al., 2012).

## Author contributions

V.A.S.J., M.B., E.A.H. and A.G. designed the experiments. V.A.S.J. and M.B. performed the experiments. V.A.S.J., M.B., E.A.H. and A.G. wrote the manuscript. All authors reviewed and approved the manuscript.

## Acknowledgments

We thank Natascha Bechtoldt, Marie Jacobovitz, and Gideon Bergheim for technical help, and Steffen Lemke, Thomas Holstein, Mark Lommel, Suat Özbek, Jochen Wittbrodt, and Aissam Ikmi for advice, comments, and sharing reagents and equipment. We also thank Christian Ackermann at the Heidelberg Nikon Imaging Center for technical support with microscopy.

